# Spatiotemporal profiling of a dominant coral’s photo-endosymbiotic assemblage indicates that acclimation is supported by phenotypic plasticity of single cells

**DOI:** 10.1101/2022.12.24.520888

**Authors:** CJ Anthony, C Lock, BM Taylor, B Bentlage

## Abstract

Coral-associated dinoflagellates (Symbiodiniaceae) are photosynthetic endosymbionts that influence coral acclimation and adaptation, as indicated by photo-physiological plasticity (phenotypic variance) in response to environmental change. Symbiont shuffling (shifts in endosymbiont community composition), changes in endosymbiont cell density, and phenotypic plasticity have all been proposed as mechanisms to adjust to environmental change. However, few studies have been able to partition which of the three strategies were responsible for observed phenotypic variance. Therefore, we quantified the biodiversity, cell density, and phenotypic variance of single cells for *Acropora pulchra-*associated Symbiodiniaceae assemblages. Using a combination of metabarcoding and flow cytometry, we simultaneously characterized Symbiodiniaceae assemblages at the community (biodiversity), population (cell density), and individual level (phenotype) under natural environmental conditions to determine whether phenotypic variation of Symbiodiniaceae communities is concomitant with either symbiont shuffling, changes in cell density, or phenotypic plasticity. Symbiodiniaceae assemblages displayed season-specific phenotypic variance, while biodiversity was geographically structured and cell density showed limited data structure. Based on these patterns, we reveal that phenotypic plasticity of individual Symbiodiniaceae cells is the source of a phenotypic variation, thus indicating that phenotypic plasticity is a mechanism for rapid acclimation to mild environmental change.

## 1 Introduction

Each coral colony supports its own unique assemblage of symbionts, with each symbiotic combination having the capacity to create a functionally distinct coral (Gates and Ainsworth, 2011). Symbiodiniaceae are dinoflagellates known for their endosymbiotic relationship with many marine invertebrates including Cnidaria, Mollusca, Porifera, Platyhelminthes, Foraminifera, and Ciliata (LaJeunesse et al., 2018). The ecological success of reef-building corals has been attributed to this endosymbiotic relationship with Symbiodiniaceae (Gault et al., 2021). Corals are highly dependent on Symbiodiniaceae for nutrient acquisition and effective calcification (Falkowski et al., 1984; Muscatine et al., 1984; Ezzat et al., 2017; Matthews et al., 2017), but environmental stress can cause a breakdown of this photo-endosymbiotic relationship, leading to the expulsion of Symbiodiniaceae from the host (coral bleaching) and often death (Brown, 1997). Climate change has increased the frequency and severity of coral bleaching globally (Hughes et al., 2017) and the survival of corals has been linked to differences in the ecological tolerances of Symbiodiniaceae (Thornhill et al., 2014; Parkinson et al., 2015; Howe-Kerr et al., 2020).

In the short term (within a generation), Symbiodiniaceae communities seem largely controlled by the host, while in the long term (across generations), environmental change may shift symbiont community composition (Baker et al., 2018; Camp et al., 2019; Howe-Kerr et al., 2020). If environments change, successful acclimation of corals may be caused by shifts in endosymbiont community composition, known as symbiont shuffling (Buddemeier and Fautin, 1993; Baker, 2003; Jones et al., 2008). For example, *Durusdinium* is more common than other Symbiodiniaceaeae genera in stressful environments (Fabricius et al., 2004; LaJeunesse et al., 2010) or after acute stress events (Baker et al., 2004; Berkelmans and van Oppen, 2006). However, endosymbiotic Symbiodiniaceae community composition can also be remarkably stable (Rouzé et al., 2019), indicating a high level of host-symbiont specificity, thus requiring a high acclimation potential through phenotypic plasticity to survive environmental change (Goulet, 2006). Species of Symbiodiniaceae have been experimentally shown to have varying rates of plasticity to environmental change (Mansour et al., 2018); therefore, the acclimation and adaptation strategy of a coral-associated Symbiodiniaceae assemblage is dependent on its constituent members and its environment.

In addition to adaptation through shuffling Symbiodiniaceae community composition, Symbiodiniaceae have several mechanisms for acclimation through phenotypic plasticity. Generally, Symbiodiniaceae acclimation revolves around modifying the efficiency and productivity of their photosystem, as it is the most direct way to regulate ATP and NADPH formation, and in turn, the quantity of harmful and beneficial metabolic byproducts (Oakley et al., 2014). Photosystem acclimation is typically tied to the modification of photosystem photochemistry (Warner et al., 1996; Ulstrup et al., 2008; Nitschke et al., 2018). The physical reorganization and regulation of photopigments or adjustments to cell morphology can also mitigate stress and promote successful acclimation to different conditions and may be a better representation of long term (months versus days) system regulation (Johnsen et al., 1994; Sawall et al., 2014; Xiang et al., 2015; Oliveira et al., 2022).

Photoacclimation *in situ* is well studied during seasonal change (Warner et al., 2002; Ulstrup et al., 2008; Sawall et al., 2014) and along light attenuation gradients including depth (Iglesias-Prieto et al., 2004; Frade et al., 2008; Lesser et al., 2010; Cooper et al., 2011) and turbidity (Hennige et al., 2008; Suggett et al., 2012). This research typically attributes observed photoacclimation patterns to (1) changes in community composition, (2) endosymbiotic cell densities, and (3) phenotypic plasticity. However, research often relies on pulse-amplitude-modulated (PAM) fluorometry (Warner et al., 1996), multi-spectral fluorometry (Hoadley et al., 2023), or high performance liquid chromatography (HPLC) (Mantoura and Llewellyn, 1983) to characterize the state of endosymbiont photosystems. These methods can be standardized to ‘phenotype’ coral holobionts (Voolstra et al., 2020), but cannot provide insight into phenotypic variance of individual symbiont cells. Alternatively, flow cytometry is an underutilized methodology that can rapidly quantify symbiont cells (Krediet et al., 2015) and generate phenotypic profiles on a per cell basis, thus providing the resolution required to identify the source of phenotypic variation (Apprill et al., 2007; Anthony et al., 2023).

Symbiont shuffling, cell density regulation, and phenotypic plasticity have all been proposed as mechanisms for Symbiodiniaceae to adjust to environmental change (Jones and Yellowlees, 1997; Baker, 2003; Baker et al., 2004; Goulet, 2006). Despite the plethora of knowledge on Symbiodiniaceae photophysiology, much of our knowledge has not closed the gap in resolving how change in phenotypic variation is explained by adaptation (shifts of Symbiodiniaceae community composition) versus acclimation (phenotypic plasticity of the existing community). Using a combination of DNA metabarcoding and flow cytometry, we simultaneously characterized the *Acropora pulchra-*associated Symbiodiniaceae assemblage at the community (biodiversity), population (cell density), and individual level (phenotype) under natural environmental conditions to understand whether phenotypic variation of Symbiodiniaceae communities is caused by symbiont shuffling, changes in cell density, or phenotypic plasticity.

## 2 Materials and Methods

### 2.1 Coral colonies

To identify the dynamics of Symbiodiniaceae assemblages under natural seasonal fluctuation on an island-wide scale, four thickets of *A. pulchra* from five reef flats (20 thickets total) were GPS-tagged around the island of Guam: Urunao (N 13.63672° E 144.84527°), West Agaña (N 13.47993° E 144.74278°), Luminao (N 13.46584° E 144.64496°), Togcha (N 13.36865° E 144.774967°), and Cocos Lagoon (N 13.24596° E 144.68475°) (Figure 1A-F).

**Figure 1.**
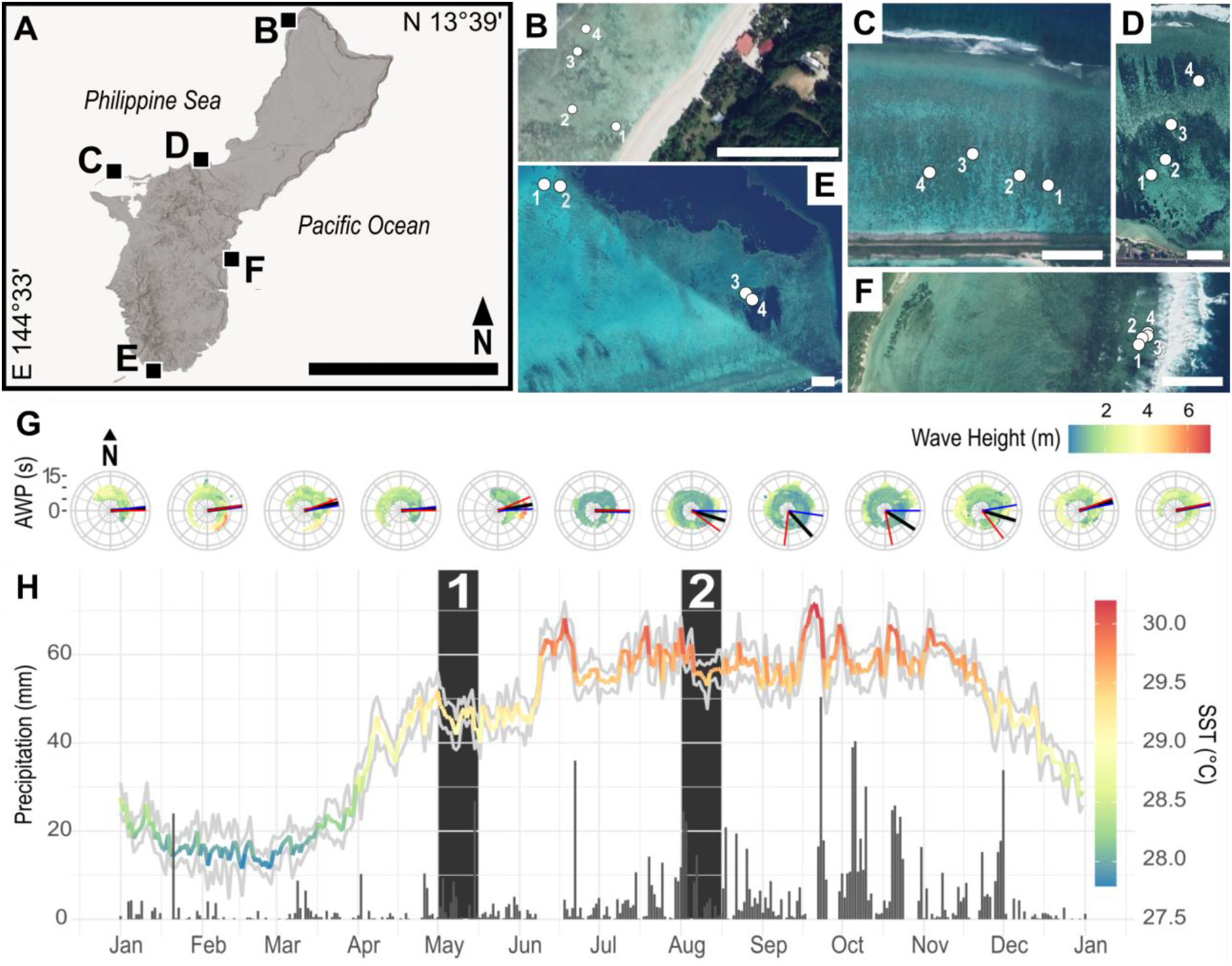
**A**) Sampling sites in Guam (scale: 20 km) with each square indicating the location of a reef flat sampled for this study: **B**) Urunao (North), **C**) West Agaña (Northwest), **D**) Luminao (West), **E**) Cocos Lagoon (South), and **F**) Togcha (East). **B-F**) Within each site, four thickets (**1-4**) of *A. pulchra* were georeferenced for repeated sampling. Images provided by Google Earth Pro v7.3.4.8248 (scale: 100 m). **G**) Spectral polar plots of aggregated historical wave data from Ritidian (**red lines**) and Ipan (**blue lines**) wave buoys. Monthly mean wave direction (**black lines**) indicated prevailing swells from the East, the windward side of Guam. (Provided by PacIOOS; www.pacioos.org). **H**) Average sea surface temperature (SST) (**spectral line**) and precipitation (**grey bars**) for 2021 showed distinct seasonal patterns. (Provided by NOAA Coral Reef Watch; www.coralreefwatch.noaa.gov) The first set of samples was collected in the first two weeks of May (**1**) during the transitional warming period, while the second set of samples was collected in the first two weeks of August (**2**) during the hot, rainy season.

### 2.2 Tissue sampling

From each site (Figure 1A-F), three tissue samples per thicket were collected by cutting two-centimeter-long pieces from at least three centimeters below the axial growth tip. This was repeated for all 20 GPS-tagged colonies during two time periods: (1) 30 April - 18 May and (2) 28 July - 15 August 2021. All samples were immediately flash-frozen in liquid nitrogen on site, then stored at -80 °C until processing. These samples were subsequently used to quantify Symbiodiniaceae biodiversity, cell density, and phenotype, deriving all variables from the same source fragment.

### 2.3 Symbiodiniaceae cell density

Coral tissue was airbrushed from the skeleton with filtered seawater (FSW) and homogenized using a vortexer followed by syringe needle-shearing, and then processed using the protocol described in Anthony et al. (2023).

Absolute cell counts were obtained by multiplying cytometry-generated cell concentrations with each sample’s dilution factor and tissue homogenate volume to determine total cell count for each coral tissue fragment. Cell density per cm2 (Equation 1) was obtained by the normalization of flow cytometry-derived cell counts to the source fragment’s skeletal surface area. To determine skeletal surface area, a three-dimensional model was created for each coral fragment built from point clouds with 0.010 mm point spacing generated by a jewelry scanner (D3D-s, Vyshneve, Ukraine). Prior to scanning, coral fragments were coated with SKD-S2 Aerosol (Magnaflux, Glenview, IL) to reduce skeletal light refraction. Point clouds of each fragment were imported into MeshLab v2020.04 (Cignoni et al., n.d.) to generate a surface mesh by Poisson surface reconstruction. Portions of the fragment that were not covered in tissue prior to airbrushing were removed from the reconstructed surface prior to surface area estimation.

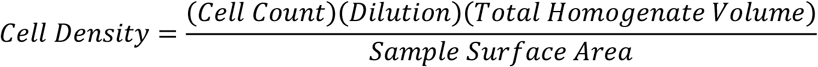

### 2.4 Symbiodiniaceae phenotyping

Three flow cytometric signatures were used to generate a phenotypic profile for Symbiodiniaceae cells: red fluorescence, forward scatter, and side scatter (Anthony et al., 2023). Red fluorescence represents relative photopigment abundance (Lesser, 1996; Lee et al., 2012; Cooper et al., 2014). Side scatter is a representation of cell shape or roughness, and forward scatter is a representation of cell size or cell volume (Mullaney et al., 1969; Steen, 1980; Shapiro, 2005; Tzur et al., 2011).

### 2.5 Symbiodiniaceae ITS2 biodiversity

Genomic DNA was extracted from tissue aliquoted prior to airbrushing (see ‘*Symbiodiniaceae cell density’* above) using a Qiagen DNeasy PowerSoil Pro Kit (Qiagen, Hilden, Germany) on a Qiacube connect liquid handling system. The ITS2 region was amplified via PCR with SYM_VAR_5.8S2 and SYM_VAR_REV primers (Hume et al., 2018) using 3 μL of DNA (10 ng/μl), 3 μl of 10 μM primer, 2.4 μl of 2.5 mM dNTP, 18.6 μl water, 3 μl buffer (10x), and 0.15 μl Taq (TaKaRa Taq™ DNA Polymerase 1U, Takara Bio USA, Ann Arbor, MI). The PCR profile included 26 cycles of 95°C for 40 s, 59°C for 120 s, 72°C for 60 s, and a final extension at 72°C for 420 s. ITS2 amplicons were multiplexed and sequenced on a NovaSeq 6000 (Illumina, San Diego, CA, USA) to generate 250 bp paired-end reads.

### 2.6 Statistical analysis

Flow cytometric data (cell density, red fluorescence, forward scatter, and side scatter) violated the assumption of parametric tests of a normal data distribution, as determined by Shapiro-Wilk tests (p < 0.001); therefore all statistical tests were non-parametric and did not assume normality.

To identify possible correlations between cell density, red fluorescence, side scatter, and forward scatter, non-parametric Spearman correlations were calculated for two groupings of paired dependent variables: (1) cell density, red fluorescence, side scatter, and forward scatter averaged to the fragment-level replicate and (2) red fluorescence, side scatter, and forward scatter without any value summarization (single symbiont cells), given that flow cytometry automatically produces paired data for each cell detected (Anthony et al. 2023). Calculations were completed using rstatix v0.7.1 (Kassambara, 2022).

Repeated measures, univariate analyses of variance (RM-ANOVA) were performed to quantify the factorial contribution of time, site, and plot to the data structure of cell density, red fluorescence, forward scatter, and side scatter using the MANOVA.RM package v0.5.3 (Friedrich et al., 2022). Main and interaction effects were resampled with 1000 non-parametric bootstrap replicates and corrected p-values were calculated for type statistics. This test neither assumes multivariate normality nor covariance matrix specificity, making it robust to repeated measure designs with factorial nesting (Friedrich et al., 2019). Prior to any statistical tests, 1000 observations for each group were randomly sampled for each cytometry replicate (often from a sample size of > 100,000 per cell measurements) to reduce the computational requirements of statistical tests and decrease the likelihood of overinterpretation.

In addition to the repeated measures tests, data from the 40 sampling events (five sites, two timepoints, and 20 colonies) were evaluated with non-parametric Kruskal-Wallis tests and pairwise Dunn’s tests, as integrated in the FSA package v0.9.3 (Ogle et al., 2022). Statistical outputs were then converted to statistical groups using _rcompanion v2.4.21 (Mangiafico, 2023). Each sample distribution was assigned to a statistical group (a-s) based on the results of pairwise Dunn’s tests, which better visualized data structure and similarity, thus improving the interpretation of previously calculated repeated measures tests. A distribution would be in the same group as another distribution if they were not statistically different (e.g., p > 0.001), while distributions would appear in different groups if they were statistically different (e.g., p < 0.001). Cell density data distributions were grouped with a standard threshold of p = 0.05, while phenotypic measurements were grouped with a threshold of p = 0.001 to avoid the overinterpretation of effect sizes made stronger by a high sample size (n = 1000) (c.f. Anthony et al. 2023).

Symbiodiniaceae community structure was compared across sites (North, Northwest, West, South, East) and seasons (May, August). After metabarcoding data was preprocessed with the SymPortal pipeline (Hume et al., 2019) and subsequently deposited on www.symportal.org (database name: 20220419_GuamWild_bentlage), relative ITS2 sequence abundances and type profiles were normalized and visualized as per Eckert et al. (Eckert et al., 2020). Multivariate homogeneity of dispersion (PERMDISP), pairwise permutation tests, and a permutational multivariate analysis of variance (PERMANOVA) were conducted on normalized ITS2 type profiles using Vegan v2.5-7 (Oksanen et al., 2019) and pairwise Adonis v0.4 (Martinez Arbizu, 2017) packages. PERMDISP and permutation tests used Bray-Curtis dissimilarity. Permutation tests were run with 9999 replicates.

To test whether underlying phenotypic variation was caused by plasticity or biodiversity, ITS2 type profiles identified by metabarcoding were mapped to their fragment’s associated phenotypic profile generated by flow cytometry. Phenotypic measurements for the North (the only site with heterogeneous Symbiodiniaceae biodiversity) were compared using a combination of Kruskal-Wallis and Dunn’s tests; with the same process described in the previous paragraph. All data curation and statistical analyses were completed with R v4.1.2 in RStudio v1.3.1073. Figures were generated and modified with a combination of ggplot2 v3.3.5 (Wickham, 2016) and InkScape v1.1 (https://inkscape.org).

## 3 Results

### 3.1 Environment

Guam’s windward (East) and leeward (West) sides are characterized by a large disparity in average wave energy. Wave energy on Guam was highest from December to March with waves, on average, coming from the East year-round (Figure 1G). In 2021, Guam did not enter a formal coral bleaching warning (Liu et al., 2018; Skirving et al., 2020) nor was bleaching observed or reported locally. Water temperatures increased steadily from March to June, remaining stable during the following four months. Precipitation followed a similar trend (Figure 1H). May represented a seasonal transition with warming waters and decreasing wave energy; August was characterized by high water temperatures and low wave energy (Figure 1G-H).

### 3.2 Symbiodiniaceae cell density

Cell density only varied with time (t = 20.81, p = 0.042), which was primarily driven by increased densities in South colonies during August (Figure 2A). All other factorial contributions were insignificant (Site: t = 0.917, p = 0.44; Time:Site: t = 2.99, p = 0.159; Site:Plot: t = 0.758, 0.516; Time:Site:Plot: t = 0.711; 0.528) (Table S1). Pairwise Dunn’s tests did not reveal any obvious factorial structuring (Figure S1A). Cell density was not correlated to any phenotypic metric (RED: ρ = 0.096, p = 0.299; FSC: ρ = 0.021, p = 0.822; SSC: ρ = 0.03, p = 0.742).

**Figure 2.**
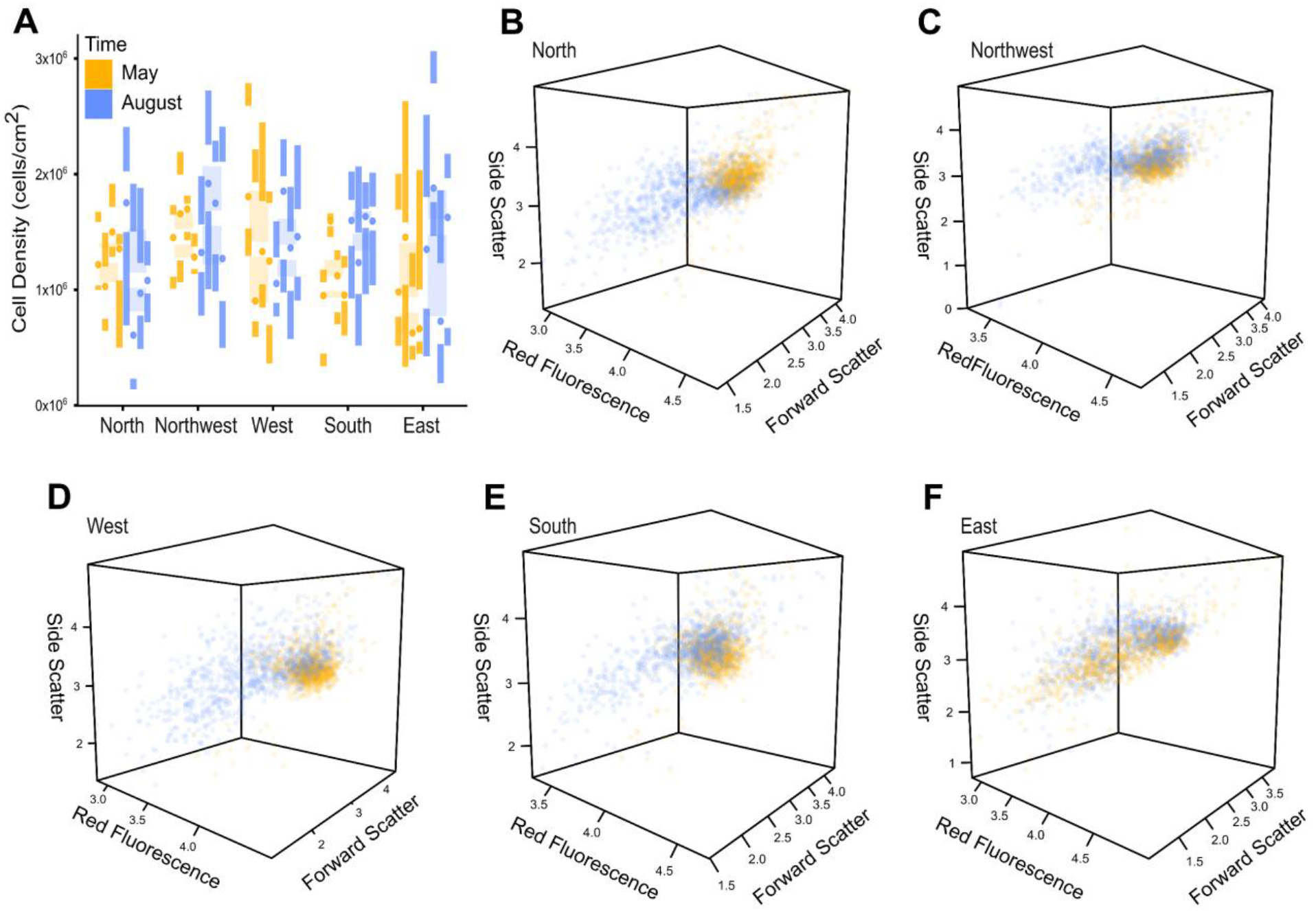
Cell density and phenotypic variance plots colored for repeated temporal sampling (May & August). **A**) Cell density illustrated no pattern, as visualized by Tufte’s box plots summarized to the plot level within each site and colored across temporal sampling points. **B-F**) Phenotypic variance illustrates a temporal pattern, as visualized by three-dimensional dot plots for North (**B**), Northwest (**C**), West (**D**), South (**E**), and East (**F**) Symbiodiniaceae assemblages.

### 3.3 Symbiodiniaceae phenotypic variation

Symbiodiniaceae phenotypic metrics were heavily influenced by time (t = 2165.490, p < 0.001), site (t = 4859.680, p < 0.001), and site within time (t = 18.661, p < 0.001) (Table S2). Time was especially influential for cell phenotype with low phenotypic variance in May and a wide phenotypic variance in August (Figure 2B-E). The East site displayed a wide phenotypic variance at both sampling time points (Figure 2F). All phenotypic variables were correlated to each other (RED-FSC: ρ = 0.38, p < 0.001; RED-SSC: ρ = 0.37, p < 0.001; FSC-SSC: ρ = 0.54, p < 0.001).

Red fluorescence (photopigment abundance) was most influenced by site (t = 291.668, p < 0.001); although it was also influenced by time (t = 1056.040; p = 0.002) (Table S2). Generally, red fluorescence declined from May to August, aside from a few plot-specific scenarios, while sites showed a more complex, case-specific partitioning of statistical groups (Figure S1B). Side scatter (cell roughness) displayed similar trends of case-specific partitioning but was especially influenced by site within time (t = 86.503, p < 0.001) and did not show any obvious large-scale pattern (Figure S1C). Forward scatter (cell size), by contrast, showed strong structuring with Site within Time (t = 322.342, p < 0.001), and displayed comparatively little within site data variation (Table S2; Figure S1).

### 3.4 Symbiodiniaceae biodiversity

Symbiodiniaceae communities of *A. pulchra* were largely dominated by *Cladocopium* C40 (Figure 3A). ITS2 type biodiversity and beta-diversity dispersion was determined by site (F = 88.793, p = 0.001; F = 11.725, p < 0.001) and not by time (F = 0.437, p = 0.761; F = 0.014, p = 0.906) (Figure 3; Table S3). ITS2 type profiles showed Symbiodiniaceae community overlap along Guam’s western coast, while southern and eastern IT2 type profiles were distinct (Figure 3).

**Figure 3.**
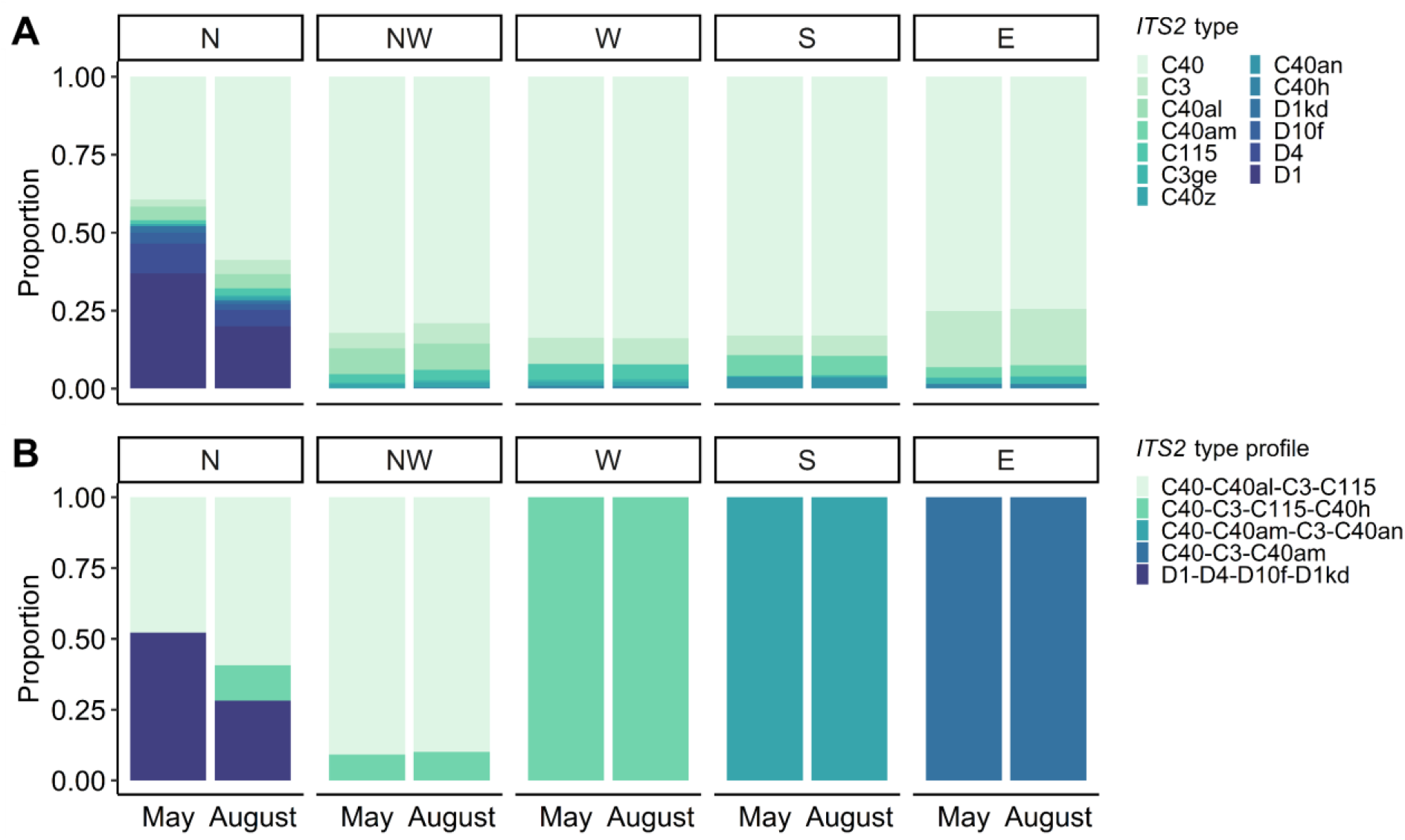
ITS2 type diversity (**A**) and ITS2 type profiles (**B**) from spatiotemporal sampling across five sites (North, Northwest, West, South, and East) and two timepoints (May and August). not statistically differentiated between sampling time points (F = 0.543, p = 0.494); however, plot-specific data suggested *Cladocopium*-*Durusdinium* partitioning from nearshore to farshore colonies, with *Durusdinium* being more common nearshore (Figure 4).

Pairwise permutation tests revealed North as an outlier, the only site with a *Durusdinium* ITS2 type profile. A pairwise permutation test of the North site across time points indicated that communities were

### 3.5 Biodiversity-phenotype association

As discussed in previous sections, only the North site showed co-dominance of *Cladocopium* and *Durusdinium* ITS2 type profiles (Figure 3). Within this site, type profiles changed from nearshore *Durusdinium-*dominated colonies to farshore *Cladocopium*-dominated colonies (Figure 4); therefore, plots were compared to evaluate whether ITS2-type profiles were associated with phenotypic variance (Figure 4). Independent from plot, type profiles did not differ in cell density (X2 = 5.760, df = 2, p = 0.056), despite a lower mean cell density in *Durusdinium*-dominated (D1-D4-D10f) colonies (Figure 4A). Plot-specific measurements illustrated a general increase in cell density from nearshore to farshore corals (Figure 4B), although accompanied by high variance. Within the North site, neighboring colonies with different ITS2 types (*Durusdinium* vs. *Cladocopium*) displayed different phenotypic profiles (e.g. colonies in Plots 2 and 3) (Figure 4C-E); however, neighboring colonies of the same ITS2 types (*Cladocopium* vs. *Cladocopium* or *Durusdinium* vs. *Durusdinium*) also demonstrated phenotypic differences (e.g. colonies in Plots 1, 3, and 4) (Figure 4C-E).

**Figure 4.**
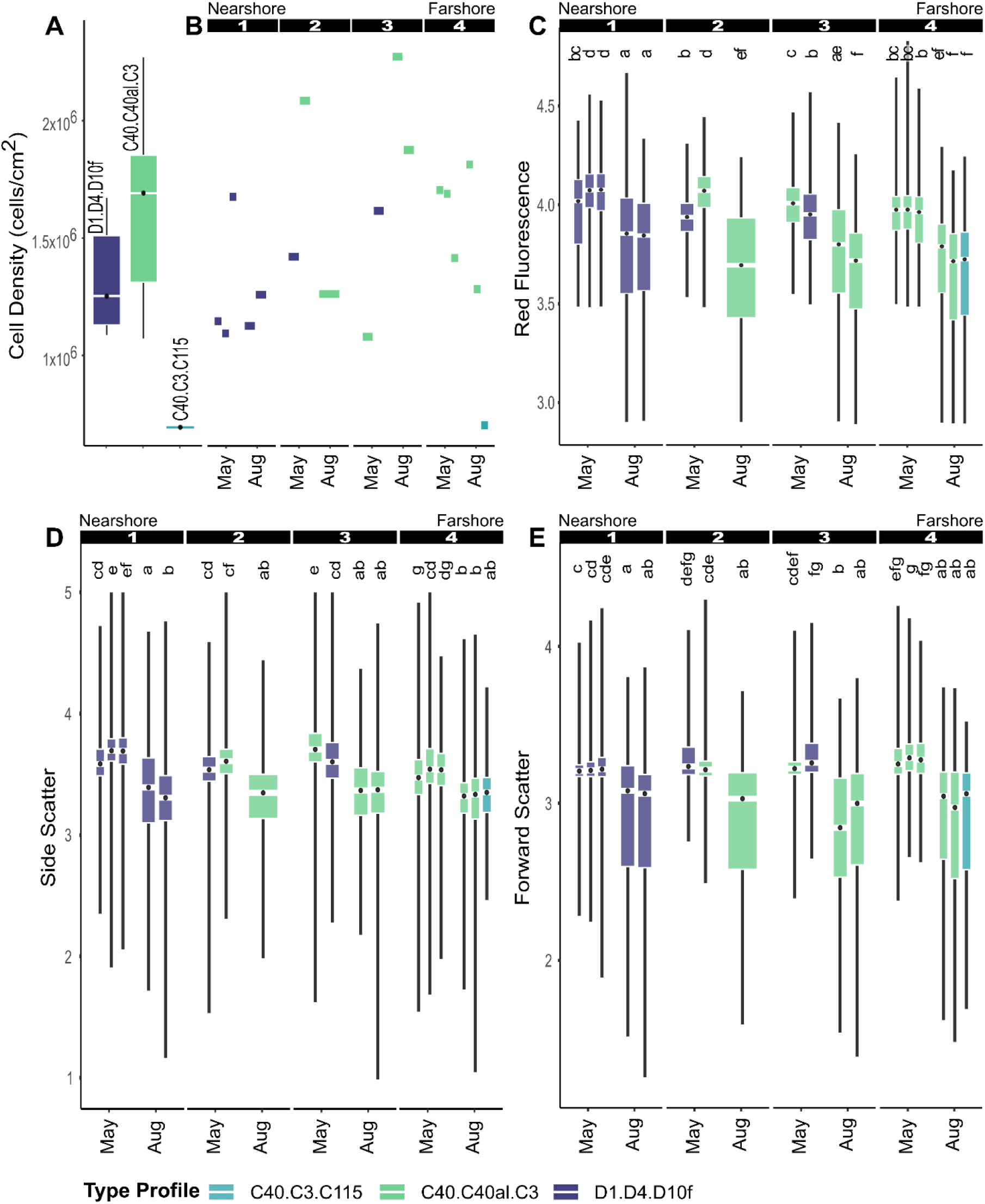
Cell density (**A-B**) and phenotypic measurements (**C-E**) were mapped to samples from the North site with known ITS2 type profiles. Sampled plots spanned across the reef flat from nearshore to farshore (1-4) (Figure 1B). Box plots (**C-E**) correspond to different colonies sampled within their respective plots (Figure 1). Letters above each boxplot (**C-E**) indicate statistical groupings (p <0.001).

## 4. Discussion

Symbiont shuffling, cell density regulation, and phenotypic plasticity have all been proposed as mechanisms for Symbiodiniaceae assemblages to adjust to environmental change (Jones and Yellowlees, 1997; Baker, 2003; Baker et al., 2004; Goulet, 2006). No trends were apparent in the cell density data (Figure 2A); however, paling and darkening between sampling time points was apparent during field sampling (Figure S2). Therefore, the visible change must have been caused by changes in Symbiodiniaceae phenotype, either by symbiont shuffling (switching to different symbiont clades) or by phenotypic plasticity (changes in morphology or physiology of cells). This hypothesis is consistent with the spatiotemporal structuring of phenotypic variance (Figure 2B-F; Table S2; Figure S1). Spatiotemporally structured phenotypic variation (Figure 2B-F), geographic structuring of ITS2 biodiversity (Figure 3), and lack of cell density patterns (Figure 2A), suggests that phenotypic plasticity of Symbiodiniaceae cells was responsible for temporal phenotypic variation.

We found that *A. pulchra*-associated Symbiodiniaceae ITS2 types were geographically structured and temporally stable (Figure 3). Long-term monitoring of Symbiodiniaceae communities found high stability and colony-level specificity in other regions (Rouzé et al., 2019); however, this high fidelity in *A. pulchra* is surprising. In *Acropora*, Symbiodiniaceae are acquired from the environment (Baird et al., 2009), suggesting a more flexible and diverse assemblage compared to other coral genera (Rouzé et al., 2017; Qin et al., 2019). Coral colonies sampled here were almost exclusively dominated by *Cladocopium* C40, with only one site containing *Durusdinium* D1 (Figure 3). Both C40 and D1 represent important lineages associated with reduced coral bleaching rates and increased coral survival following stress (Jones et al., 2008; Mieog et al., 2009; Rouzé et al., 2017; Qin et al., 2019).

Species of Symbiodiniaceae have been experimentally shown to have varying rates of plasticity to respond to environmental change (Mansour et al., 2018); therefore, we expected that phenotypic variation of a coral-associated Symbiodiniaceae assemblage would differ between different Symbiodiniaceae assemblages. Only the North site provided any insight to this hypothesis given the codominance of two ITS2 type profiles (Figure 4). Colonies from nearshore (*Durusdinium* D1-dominated) to farshore (*Cladocopium* C40-dominated) displayed some mild differences in phenotypic characteristics, such as the higher abundance of photopigments (high red fluorescence) in nearshore colonies (Figure 4C). However, the phenotypic variance between neighboring *Durusdinium* and *Cladocopium*-dominated colonies (e.g. colonies in Plots 2 and 3) was no different than neighboring colonies of the same ITS2 types (e.g. colonies in Plots 1, 3, and 4) (Figure 4). Again, the data indicates that phenotypic variation is most likely caused by phenotypic plasticity of individual Symbiodiniaceae cells.

In general, the dominance of thermotolerant Symbiodiniaceae, *Cladocopium* C40 and *Durusdinium* D1, in *A. pulchra* on Guam’s reef flats may be the result of selection, following mass coral mortality caused by multiple bleaching events over the last decade (Raymundo et al., 2017, 2019). The nearshore to farshore partitioning of *Durusdinium* D1 to *Cladocoium* C40 dominated *A. pulchra* colonies may be a fine scale adaptation to long term chronic stressors or environmental conditions. Under such extreme environmental stress successful acclimation may be caused by symbiont shuffling (Buddemeier and Fautin, 1993; Baker, 2003; Jones et al., 2008; Zhu et al., 2022). Perhaps *Durusdinium* was selected for its higher tolerance to warmer nearshore waters (Stat et al., 2008; Keshavmurthy et al., 2014; Silverstein et al., 2017).

Symbiodiniaceae communities of Guam’s *A. pulchra* populations did not display changes in cell densities, or shuffling of communities, but rather showed signs of seasonal environmental acclimation through phenotypic plasticity. These results suggest that Symbiodiniaceae phenotypic plasticity of individual cells is the primary mode of acclimation to environmental change across seasons (Discussed further in File S1). Changes in cell density are more likely an acute response to extreme stress, such as coral bleaching, while symbiont shuffling is likely a mechanism for long-term adaptation after acute stress events that select for specific Symbiodiniaceae clades. Using the same framework presented here, characterizing Symbiodiniaceae phenotypic variation with higher temporal resolution and samples from extreme events, has the potential to provide important insights into the dynamics and limits of Symbiodiniaceae acclimation *in situ*.

## 5 Data Availability Statement

All data is publicly available. Cell density and phenotypic profiling data, along with corresponding original code has been deposited on GitHub (Permanent doi will be released prior to publication; https://github.com/AnthonyCuog/SpatiotemporalPhenotypicProfiling). ITS2 metabarcoding data is publicly available at www.symportal.org (database name: 20220419_GuamWild_bentlage).

## 6 Author contributions

CJA: conceptualization, investigation, methodology, data curation, formal analysis, visualization, writing–original draft, writing – review and editing. CL: methodology, data curation, formal analysis, investigation, and writing – review and editing. BMT: formal analysis, writing – review and editing, and supervision. BB: conceptualization, methodology, investigation, resources, writing – review and editing, supervision, and funding acquisition. All authors contributed to the article and approved the submitted version.

## 8 Acknowledgements

We wish to thank Dr. Cheryl Ames, Dr. Héloïse Rouzé, and Dr. Shinichiro Maruyama for providing their expertise and mentorship. We would also like to thank Marine Laboratory boat captain, Jonathan “Nanny” Perez, for assisting with field collections in Cocos Lagoon, and the Ritidian Eco Beach Resort for providing land access to the pristine reef flats of Urunao.

