## Supplementary Material for "Spatiotemporal profiling of a dominant coral’s photo-endosymbiotic assemblage indicates that acclimation is supported by phenotypic plasticity of single cells"

### 1 Supplementary Data

All data is publicly available. Cell density and phenotypic profiling data, along with corresponding original code has been deposited on GitHub (Permanent doi will be released prior to publication; https://github.com/AnthonyCuog/SpatiotemporalPhenotypicProfiling). ITS2 metabarcoding data is publicly available at www.symportal.org (database name: 20220419_GuamWild_bentlage).

### 2 Supplementary Figures and Tables

#### 2.1 Supplementary Tables

**Supplementary Table 1.** RM-ANOVA testing the factorial contribution to variation in cell density.

| | **POPULATION** | |
|---|---|---|
| | Cell Density | |
| | t | p |
| **Time** | 20.81 | 0.042 |
| **Site** | 0.917 | 0.44 |
| **Time:Site** | 2.99 | 0.159 |
| **Site:Plot** | 0.758 | 0.516 |
| **Time:Site:Plot** | 0.711 | 0.528 |

**Supplementary Table 2.** RM-MANOVA and three RM-ANOVAs testing the factorial contribution to variation in red fluorescence, side scatter, and forward scatter.

| | **INDIVIDUAL** | | | | | | | |
|---|---|---|---|---|---|---|---|---|
| | RED, FSC, SSC | | RED | | SSC | | FSC | |
| | t | p | t | p | t | p | t | p |
| **Time** | 2165.49 | <0.001 | 1056.0<br>4 | 0.002 | 33.978 | 0.02 | 5071.4<br>1 | 0.001 |
| **Site** | 4589.68 | <0.001 | 208.90<br>3 | <0.001 | 44.468 | 0.002 | 414.25<br>6 | <0.001 |
| **Time:Site** | 18.661 | <0.001 | 48.343 | 0.003 | 86.503 | <0.001 | 322.34<br>2 | <0.001 |
| **Site:Plot** | 162.621 | 0.116 | 2.643 | 0.172 | 2.863 | 0.17 | 2.073 | 0.230 |
| **Time:Site:Plot** | 93.414 | 0.242 | 0.808 | 0.489 | 2.434 | 0.195 | 1.567 | 0.313 |

**Supplementary Table 3.** Outputs of PERMANOVA and PERMDISP to determine factorial contributions for ITS2-type diversity and ITS2 beta-diversity, respectively.

| | **COMMUNITY** | | | |
|---|---|---|---|---|
| | **ITS2-type** | | **Beta-diversity** | |
| | **F** | **p** | **F** | **p** |
| **Time** | 0.437 | 0.761 | 0.014 | 0.906 |
| **Site** | 88.793 | 0.001 | 11.725 | <0.001 |

### 2.2 Supplementary Figures

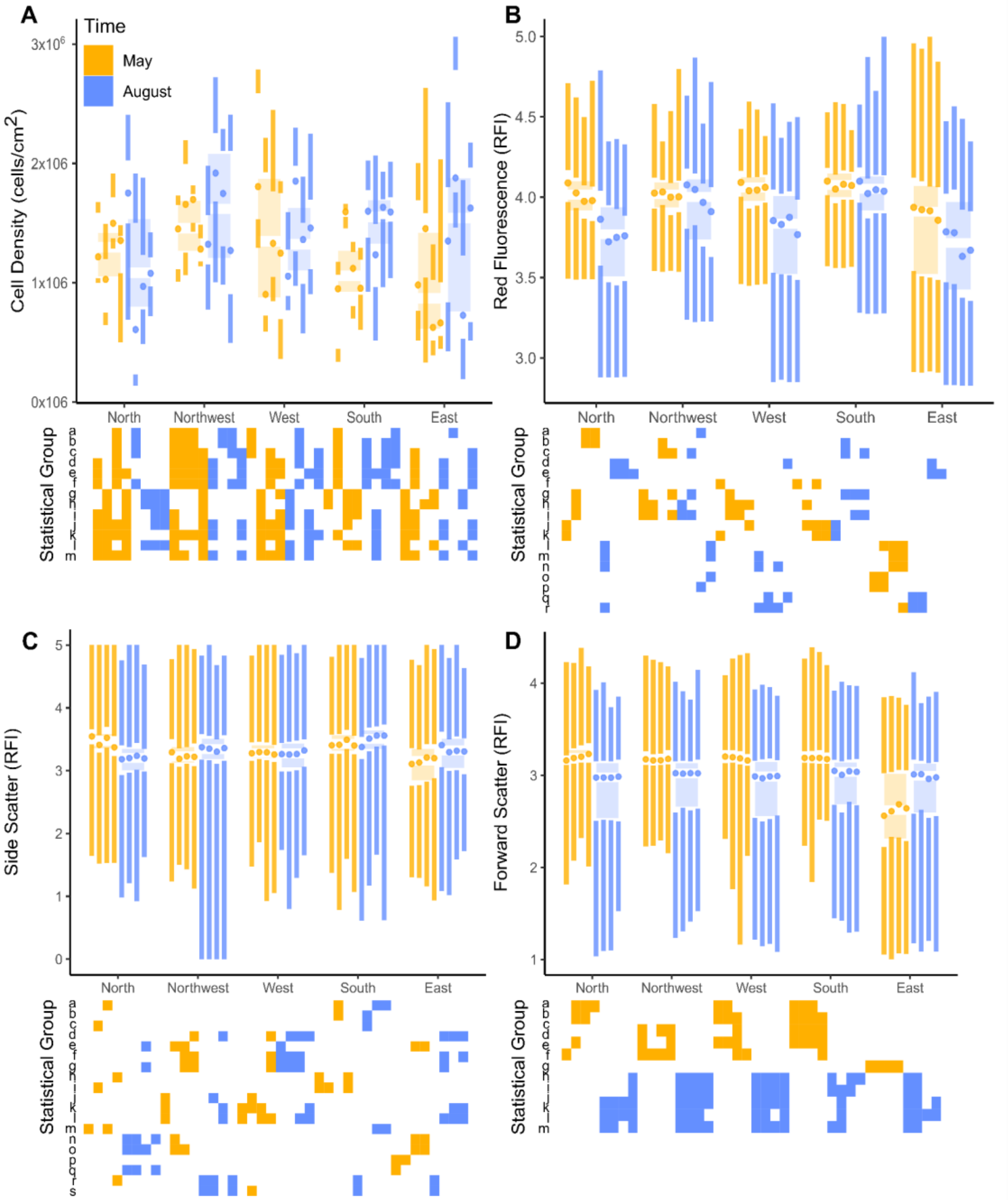


**Supplementary Figure 1**. Cell density and phenotype variance visualized as boxplots (Site * Season) and Tuft's boxplots (Plot * Site * Season). Colors indicate temporal sampling. Statistical groupings (squares) supported by pairwise Dunn's tests (**A:** $p < 0.05$; **B-D:** $p<0.001$) reveal data structure hidden by most visualizations and broader statistics. **A**) Cell density shows almost no statistical structure. **B-C)** Red fluorescence (photopigment abundance) and side scatter (cell roughness) show statistically supported, case-specific factorial contributions. **D)** Forward scatter shows high temporally structured statistical groupings.

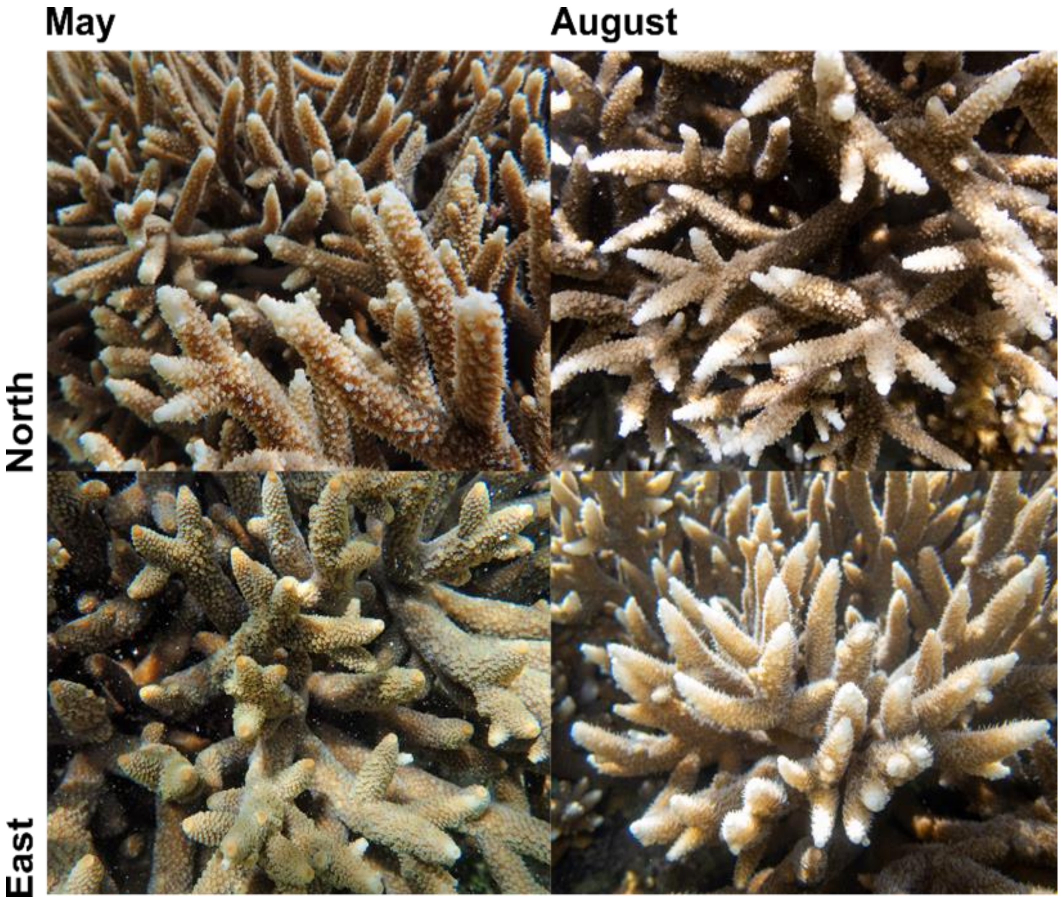


**Supplementary Figure 2**. Colonies in different sites demonstrated different levels of pigmentation not only from each other, but also across seasons, as demonstrated by repeated photographs during May and August of the same colonies from the North and East sites.

### 2 Supplemental Discussion

#### 2.1 Dynamics of Phenotypic Traits

We felt the following discussion was outside the scope of our main manuscript; however, we still find it important to discuss and present a series of hypothesize related to the phenotypic metrics derived from flow cytometric profiling. The following discussion is in place to catalyze discussion and future research.

Red fluorescence decreased from May to August in all colonies (Figure S1) indicating an overall reduction of photopigments; the generally expected result based on other literature (Porter et al., 1997; Brown et al., 1999; Mass et al., 2007); however, an increase in photopigments to seasonal warming has also been reported (Sawall et al., 2014). Cell roughness, measured by fluorescent side scatter, illustrated a very similar data structure (Figure S1C). While poorly studied, cell roughness may be another method for cells to control their light intake, and in turn their productivity. Xiang et al. (Xiang et al., 2015) found that in lower light conditions, Symbiodiniaceae reduced the amount of surface adhesion proteins produced, and gained a smoother, less complex surface morphology. Perhaps rougher cells scatter a greater amount of light, thus limiting the amount of light available to intercellular photosynthetic plastids while a smoother cell could scatter less light, allowing more access for light to reach the light harvesting structures. Unfortunately, the lack of clear spatiotemporal structuring in our side scatter data makes it difficult to resolve any relationship between side scatter and environment.

In contrast to photopigment fluorescence and cell roughness, cell size had an extremely clean temporal data structure, primarily demonstrating larger cells in May and smaller cells in August (Figure S1D). Perhaps this pattern is a broad, seasonal pattern related to the rate of cell division. Under heat stress, Symbiodiniaceae proliferation decreases and cells swell (Fujise et al., 2018), which would lead to us detecting larger cells; this is not what we saw as cells were smaller during the warm season in all except one site. While we do not see signs of heat stress, warmer temperatures do slow Symbiodiniaceae cell growth and decrease photosystem productivity (Karim et al., 2015). This is more representative of the pattern observed here; however, one site complicates this interpretation.

East colonies demonstrated a different phenotypic profile than the other sites with a much higher phenotypic variance in May than other sites (Figure 2). In May, windward colonies (East) displayed a reciprocal pattern in cell size compared to other sites, starting with smaller cells in May and converging on the cell sizes more characteristic of the rest of the island in August. Despite this, the temporal change in red fluorescence and side scatter of East corals was not much different compared to other sites (Figure S1B-C). The only known surviving *A. pulchra* on the windward side of Guam occurs in a limited area near the East site's algal ridge (Figure 1F). From December to May, the windward side experienced a high frequency of large swells (Figure 1G), which likely improved gas exchange (Finelli et al., 2006) and increased the availability of inorganic carbon, conditions that would increase the photosynthetic rates of Symbiodiniaceae (Dennison and Barnes, 1988), and in turn the abundance of metabolic byproducts and cellular population. Interestingly, these windward coral colonies displayed a distinctive green hue in May, presumably caused by green fluorescent protein (GFP), which was absent in August and not observed at other sites (Figure S2). If the coral host does not regulate its Symbiodiniaceae endosymbionts, either through ROS scavenging or cell density regulation, the relationship can switch from symbiotic to parasitic (Cunning and Baker, 2012; Morris et al., 2019). Coral-host GFP expression has been linked to stress mitigation through shading of photosymbionts (Lyndby et al., 2016) and antioxidant activity (Palmer et al., 2009). Perhaps the visible expression of GFP (Figure S2), coinciding with high phenotypic variation (Figure 2), smaller cells (Figure S1D), and a normal cell density (Figure 2A), was a form of host regulation to avoid a breakdown of symbiosis. In June, warming waters, decreased water flow, and increased rainfall may have made host-mediated symbiont regulation unnecessary amidst La Niña, which could explain the island-wide convergence of phenotypic variance (Figure 2, S1). This is simply a hypothesis and requires validation, but we can at least conclude that colonies in the east site were under different environmental pressures, leading to distinct phenotypic patterns and acclimation strategies.
